# Data-Driven Decomposition of Precisely Aligned Fiber-Orientation-Preserving Probabilistic Tractography Connectomes with Non-Negative Matrix Factorization

**DOI:** 10.64898/2026.11.08.765179

**Authors:** Alexander Zhou, Timothy S. Coalson, Burke Q. Rosen, Hossein Rafipoor, Fredrik Lange, Naofumi Yoshida, Andrea Yang, Takuya Hayashi, Stamatios N. Sotiropoulos, Saad Jbabdi, David C. Van Essen, Matthew F. Glasser

## Abstract

The major goals of the Human Connectome Project (HCP) were to characterize the structural and functional connectivity of the human brain. Although there has been tremendous work done for functional connectivity, including application of a variety of data driven decomposition approaches, structural connectivity from the HCP has seen less attention. This work presents a collection of methodological advances that, in combination, enable similar approaches to be applied to diffusion tractography data. Here, we use multi-modal non-linear volumetric alignment with FSL’s MultiMOdal Registration Framework (MMORF) to precisely align white matter tracts and globular gray matter nuclei, use the White matter Hierarchical Modelling (WHIM) method to align voxel-wise fiber orientations, enable efficient GPU-accelerated computation and storage of full resolution probabilistic gray matter x white matter x fiber sparse tractography matrices in precise alignment across individuals using the aforementioned MMORF and WHIM alignments together with areal-feature-based cortical surface registration. These tractography dense connectomes can be averaged across individuals and sparse, computationally efficient non-negative matrix factorization (NMF) can be applied to perform data-driven decompositions into white matter tracts and gray matter projection territories. The primary outputs of this effort to date are 1) A set of HCP software pipelines enabling such analyses, 2) A new unbiased HCP-derived volumetric standard space that is representative of the young adult brain and follows ACPC orientation conventions across individuals. 3) Reproducible NMF decompositions of HCP young adult tractography data into white matter tracts and gray matter projection territories, which could be projected using non-negative dual regression back to individuals. These data represent a starting point for a new generation of data-driven atlasing efforts for human white matter, and these methods can be applied to study the white matter in development, aging, and brain diseases.

## Introduction

The Human Connectome Project (HCP) aimed to characterize the connectivity of the human brain using non-invasive diffusion and functional MRI at unprecedented spatial resolution (Van Essen, Ugurbil, et al., 2012; Van Essen et al., 2013). In particular, diffusion MRI data were acquired at 1.25mm isotropic diffusion MRI data with customized hardware and acquisition protocols that pushed the limits of in vivo human imaging, specifically to advance data preprocessing and analysis of tractography (Sotiropoulos et al., 2013). Although accurately characterizing the area-to-area structural connectome with diffusion tractography has proven to be quite difficult (Van Essen et al., 2014), white matter fascicles (and their gray matter projection territories) present a potentially more tractable problem: For example, neurosurgically-relevant white matter tracts can now be reconstructed routinely in individual patients with brain tumors or epilepsy (Farquharson et al., 2013; Jeurissen et al., 2013), and population atlases of white matter tracts have also been developed.

These atlases have broadly followed two approaches. The first defines the white matter tracts and fascicles using predefined anatomical regions of interest or expert curation (Yeh et al., NeuroImage 2018) (Warrington et al., 2020); however, such atlases will only be as good as the neuroanatomical assumptions built into them. The second identifies tracts in a data-driven manner by decomposing a tractography “dense connectome” – a matrix of gray matter points by white matter voxels using data-driven algorithms like Independent Components Analysis (ICA; O’Muircheartaigh & Jbabdi, 2018) or Non-negative Matrix Factorization (NMF; Thompson et al., 2020) akin to those commonly used in functional MRI (e.g., Smith et al., 2012) and EEG (Makeig et al., 1995), though such approaches have received only a small fraction of the attention in diffusion MRI that has been given to them for functional signals. Such data-driven approaches will ideally identify neuroanatomically expected tracts, but may also result in new discoveries, both of fascicles, but also of finer-grained topographies within fascicles. NMF has an advantage over ICA in that tractography streamline counts are inherently non-negative, and thus negative lobes of ICA components are difficult to interpret. Indeed, NMF has been shown to successfully recover known white matter fascicles from gray matter x white matter tractography matrices (Thompson et al., 2020) and also to produce the territories of gray matter to which these fascicles project (i.e., gray matter projection territories).

However, these prior decompositions operated on two-dimensional gray matter × white matter matrices, which record how many streamlines connect each gray matter location to each white matter voxel, but not which fiber population within that voxel the streamlines followed. This approach is a substantial simplification of the available data, because high angular resolution diffusion imaging data, like that acquired by the HCP, reveals up to three fiber orientations in most 1.25mm isotropic imaging voxels (Sotiropoulos et al., 2013). For example, within the white matter of the centrum semi-ovale, there are three-way crossings of axons from the corpus callosum (running medially to laterally), the cortico-spinal tract (running superior to inferiorly), and the superior longitudinal fasciculus (running anterior to posteriorly). A gray matter × white matter × fiber matrix that retains the fiber orientation chosen by each streamline avoids this problem, and FSL’s probabilistic tractography has been able to record such orientations since early in the HCP. In practice, however, whole human brain application has been infeasible for three reasons. 1) The implementation of the gray matter x white matter x fibers counting matrix was limited to a single CPU thread. 2) Such a storage format, even in a sparse mode, required vast amounts of memory relative to what was available at the time for a whole human brain at 1.25mm isotropic diffusion resolution. 3) Prior implementations of ICA and NMF (O’Muircheartaigh & Jbabdi, 2018; Thompson et al., 2020) often operated on downsampled versions of the simpler and smaller gray matter x white matter matrices relative to the native 1.25mm resolution of HCP 3T diffusion MRI data, but such downsampling would be poorly determined if one tried to maintain a connection to the actual fiber orientations in the white matter. Thus, capabilities in both FSL’s probabilistic tractography algorithm and the HCP’s Connectome Workbench lay dormant (aside from a couple of macaque hemispheres, as reported by Donahue et al. (2016), that required over a month to produce the proof-of-concept data). That said, there have been tremendous improvements in speed and capacity of computer hardware since the HCP diffusion data were acquired.

Comparison across individuals is another important consideration. Although initial efforts used nonlinear volumetric registrations based on T1-weighted (T1w) images (O’Muircheartaigh & Jbabdi, 2018; Thompson et al., 2020), T1w images have very little contrast within the white matter. Within the cerebral cortex (Coalson et al., 2018; Robinson et al., 2014, 2018), it is well-known that multi-modal surface-based alignment outperforms folding-based alignment of cortical areas and functional networks and, thus, this motivates the use of a multi-modal nonlinear volumetric alignment to improve the alignment of white matter tracts and globular gray matter subcortical nuclei. Recently, FSL’s MultiMOdal Registration Framework (MMORF) algorithm became available, with the ability to align T1-weighted (T1w), T2-weighted (T2w), and diffusion tensor images (DTI) (Lange et al., 2024). In addition to spatially aligning white matter tracts, fiber orientations must be reoriented according to the local deformation of the warp (Alexander et al., 2001), and the multiple fiber orientations in each white matter voxel must be matched across individuals, i.e., in “fixel” space (Raffelt et al., 2015). Fortunately, the recently developed White matter Hierarchical Modelling (WHIM) tool (Rafipoor et al., 2025) addresses both issues: it reorients fiber orientations using the local Jacobian of the warp and establishes fiber correspondences by hierarchical modeling, aided by the improved interindividual alignment provided by MMORF. This approach substantially improves the interindividual alignment of gray matter x white matter x fiber matrices.

We also benefitted from advances in non-negative matrix factorization algorithms. Even if we manage to compute many precisely aligned whole human brain gray matter x white x fiber dense connectomes and average them, such a dataset would be 2.22 TB as a full matrix. Thus, maintaining all operations in sparse format, including NMF computation is highly beneficial. Although the open-source library scikit-learn has a python implementation of NMF capable of handling sparse matrices, we found it slower than an R-based implementation of sparse NMF originally designed for single-cell transcriptomic analyses (DeBruine et al., 2024). Speed is of the essence, as data-driven decomposition algorithms are ideally run multiple times, their results clustered, and the centrotypes of those clusters used to initialize a final decomposition to ensure that final decomposition is reproducible, as in ICASSO (Himberg & Hyvarinen, 2003).

Here, we combine all of the above methods and technical advances into the HCP’s pipelines using (i) MMORF to improve alignment of white matter tracts and globular gray matter nuclei; (ii) WHIM for interindividual alignment of fibers, probabilistic tractography; (iii) enabling GPU computation and sparse storage of gray matter x white matter x fibers matrices that are counted according to WHIM aligned individual fiber populations, and (iv) using sparse, reproducible non-negative matrix factorization of group average gray matter x white matter x fibers matrices into white matter fascicles and their gray matter projection territories. We provide preliminary evidence that these tools yield reproducible, high-resolution NMF decompositions of dense tractography matrices into classical white matter fascicles and their gray matter projection territories. Additionally, we produce a new young adult HCP-derived and precisely multi-modally aligned standard volume space that is both unbiased relative to the average human brain (unlike MNI; Glasser, Smith, et al., 2016), adheres to the standard AC-PC convention (unlike OMM-1; Lange et al., 2024), and is based on young adults (unlike OMM-1, which is based on older adults) and hence serves as a bridge to both developmental and aging analyses.

## Methods

### Data acquisition and preprocessing

#### Participants

The data for this study are from 957 participants in the Human Connectome Project Young Adult (HCP_YA) (Van Essen et al., 2013; Van Essen, Ugurbil, et al., 2012) having full diffusion MRI data acquired at 3T (total of 6 runs) together with T1-weighted (T1w) and T2-weighted (T2w) structural MRI and resting state fMRI data allowing them to be processed with MSMAll (*HCP-Young Adult 2025 Release*, 2025; Robinson et al., 2014, 2018) areal feature-based registration (see below). The participants were healthy young adults between 22 and 35 years of age. 41 retest subjects were rescanned across all modalities in a separate session with an interval of 18-328 days and a mean of 135 days.

#### Imaging Parameters

The acquisition was previously described in (Glasser et al., 2013; Sotiropoulos et al., 2013; Uğurbil et al., 2013). All imaging data were acquired on a customized 3 T Siemens Connectom scanner (a modified Skyra platform) equipped with a high-performance SC72 gradient coil (maximum gradient strength of 100 mT/m and slew rate of 200 T/m/s) and a 32-channel head coil. Structural MRI consisted of high-resolution 0.7 mm isotropic 3D T1-weighted MPRAGE (TR = 2400 ms, TE = 2.14 ms, TI = 1000 ms, flip angle = 8°) and 3D T2-weighted SPACE images (TR = 3200 ms, TE = 565 ms, variable flip angle), each typically acquired twice per participant. Diffusion-weighted MRI (dMRI) data were acquired in two phase reversed averages at 1.25 mm isotropic resolution with a multi-shell scheme with b-values of 1000, 2000, and 3000 s/mm², comprising 90 non-collinear gradient directions per shell (270 total unique diffusion-weighted directions) interleaved with 18 b = 0 s/mm² volumes.

#### Non-Diffusion MRI Preprocessing

All data were preprocessed using the HCP pipelines (Glasser et al., 2013) specifically using the structural preprocessing pipelines (PreFreeSurfer, FreeSurfer, and PostFreeSurfer), the functional preprocessing pipelines (fMRIVolume and fMRISurface), the multirun spatial ICA+FIX fMRI data denoising pipeline (Glasser et al., 2018, 2019; Salimi-Khorshidi et al., 2014), multi-modal “MSMAll” areal-feature-based surface registration (Glasser, Coalson, et al., 2016; Robinson et al., 2014), temporal ICA denoising (Glasser et al., 2018, 2019; Salimi-Khorshidi et al., 2014), and Transmit correction of T1w/T2w myelin maps (Glasser et al., 2022). The data used in this study is thus aligned with the HCP-YA 2025 data release.

#### Diffusion MRI Preprocessing

We elected to reprocess the HCP-YA diffusion data for this publication because of significant advances in diffusion MRI preprocessing since the HCP-YA diffusion data were initially preprocessed and released. These advances included (1) a topup configuration adapted to the 1.25 mm resolution of the HCP-YA diffusion data, with the warp resolution of the final iteration matched to the voxel size, correspondingly adjusted regularization, and no subsampling (Andersson et al., 2003); and (2) additional eddy options for outlier replacement (Andersson et al., 2016) and for correction of within-volume movement (Andersson et al., 2017) and susceptibility-by-motion interaction (Andersson et al., 2018). As such, the diffusion data in this study has been processed using similar methods to the HCA and AABC diffusion data (Harms et al., 2018).

Diffusion tensor data were derived from the preprocessed diffusion data using DTIFIT (Jenkinson et al., 2012) from FSL. However, since DTIFIT operates on a single diffusion shell, the lowest diffusion weighted shell (b<1200) was selected due to its high signal to noise ratio. Up to 3 probabilistic fiber orientations were fit with FSL’s bedpostX model=3 algorithm (Behrens et al., 2007; Jbabdi et al., 2012) using correction for gradient nonlinearities with rician noise modeling and burning period of 3000 (Hayashi et al., 2021).

### Volumetric alignment through MMORF

Accurate voxel-wise alignment across subjects within the deep white matter and globular subcortical gray matter is critical for the generation of high-quality tract atlases (and globular non-neocortical nuclear parcellations). To improve the correspondence within these structures across participants, we used FSL’s MultiModal Registration Framework (MMORF) (Lange et al., 2024) for multi-modal nonlinear registration. Compared with its predecessor FNIRT (Andersson et al., 2007), which was used during HCP preprocessing to register subjects to MNI space using only T1w images, MMORF offers multimodal registration using both scalar (T1w and T2w) and tensor (DTI) images. Allowing the simultaneous consideration of both structural and diffusion derived information into the registration process allows for superior correspondence across deep white matter tracts and globular gray matter nuclei but avoids overfitting the diffusion data, as it still follows anatomical constraints compared with single modal diffusion registrations.

**Figure 1:**
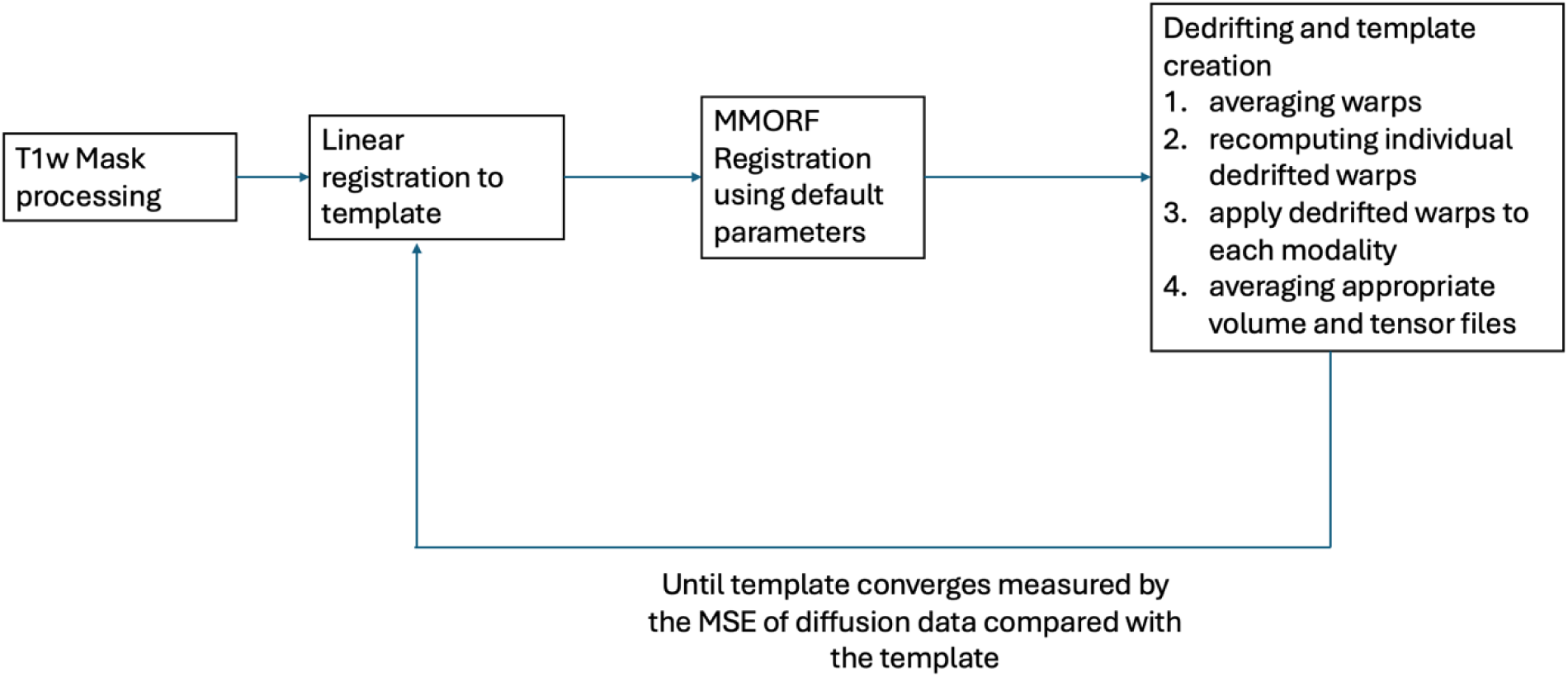
The template creation framework. Schematic summarizing the template generation process. The template generation first uses linear registration with FSL’s FLIRT followed by non-linear registration by MMORF, followed by dedrifting and averaging the dedrifted volumes and tensors to create a new template. This process is iteratively repeated until the data converges as measured by diffusion MSE relative to the prior template iteration.

#### Template generation

A registration template represents the average anatomy of a population under a given registration objective, and is therefore shaped both by the registration framework used to construct it and by the population from which it was derived. We evaluated existing templates against these two criteria before determining that a new, study-specific template was required.

The MNI template (Grabner et al., 2006) was generated previously, but is limited in two important ways. First, it does not include diffusion-derived contrasts, and second it has a substantial amount of registration drift in it (Glasser, Coalson, et al., 2016; Glasser, Smith, et al., 2016), resulting in a template brain ∼40% larger than the average volume of the original individual brains (Abdollahi et al., 2014; Van Essen, Glasser, et al., 2012). A better potential registration basis is the OMM-1 template, which is constructed using the MMORF registration framework and therefore does not share the modality limitations of the MNI template and has been appropriately dedrifted to its population. However, OMM-1 was built from UK Biobank data, with individuals aged 51–55 years at the time of the data used for OMM-1 construction. It also used T1w, FLAIR, and diffusion data, and not the T2w data available in HCP-YA. Registering our data to a template derived from a substantially older population risks introducing characteristics of the older population such as larger ventricles into the aligned data due to a mismatch between the template population and the 22–35-year-old young adult participants of the HCP-YA — a population meaningfully distinct from that used to create OMM-1. Because template-induced bias can arise independently from both (1) differences in the modalities used during template construction and (2) differences in the biological/demographic characteristics of the template population relative to the target dataset, neither the MNI Nonlinear nor the OMM-1 template was suitable: the former was limited on the modality axis and had significant baked in registration drift, and the latter was mismatched on the population axis and the modality axis to a lesser extent. Finally, OMM-1 was rigidly aligned to MNI; however, in doing this, its orientation no longer matches the expected ACPC convention of volumetric standard spaces (because of the drift in MNI space, it is difficult to accurately rigidly align OMM-1 to MNI). We elected to retain the ACPC convention in our new young adult HCP-derived template.

To retain the advantages of MMORF’s multimodal registration objective while preserving the biological characteristics specific to our population, we therefore constructed a new template using our own HCP-YA data. The process of template generation involves iterations of first registering individuals to an existing template and then doing a group wise dedrift to obtain the updated template.

#### Registration stage

To register individual subjects, several preprocessing steps need to occur to allow for the correct input into MMORF. The 8:1 ratio inside the mask and outside of the mask are maintained, consistent with the original OMM template to reduce regularization in the brain compared to skull and surrounding regions for T1w images, throughout template generation. Furthermore, an initial linear registration is done via FSL’s FLIRT based on the T1w image to the template with cubic spline interpolation to provide higher accuracy and reduced smoothness compared with the default trilinear interpolation. The nonlinear MMORF registration was conducted with T1w, T2w, and diffusion fiber orientations as the three modalities with the rest of the parameters specified by the hyperparameter search described in the original paper (Lange et al., 2024). The individual warps produced by MMORF registration are combined with the initial FLIRT linear transformation, producing the final transformation from individual space to template space.

#### Dedrifting and Template Generation stage

To generate an unbiased population template and minimize template drift and overfitting across iterations, dedrifting was performed. Because the ACPC alignment of the OMM-1 template differs from that of the MNI nonlinear space, both the (post-subject-ACPC) linear and nonlinear components of the individual transformations were included in the dedrifting procedure. Although averaging the linear and nonlinear transformations can result in a change in the overall scale of the resulting population space, the ACPC alignment largely minimizes the rotational variability prior to registration, reducing the potential for rotational differences to manifest in spatial shrinkage. Furthermore, dedrifting both the linear and nonlinear components prevents systematic translations and rotations from accumulating across template-generation iterations. Avoiding this accumulation was prioritized over preserving the global scale of the initial template, as continued drift in position and orientation would otherwise cause the template coordinate system to progressively deviate from a stable population-average space. This produces a new coordinate space representing the mean anatomical configuration of the population rather than retaining the global transformations inherited from the initial template. That said, we preserved AC-PC orientation convention in our new template.

The nonlinear warpfields for each subject are represented as a field of displacement vectors from the template image to the corresponding location in the individual image (which was already rigidly rotated and translated into approximate ACPC alignment). These vector fields were averaged across subjects to find the vector field that represents the group average drift that occurred during an iteration of template refinement. The inverse of this drift transformation was composed with each individual’s original transformation to move the image features in the next template back to their average position within the group of subjects. The resulting transformations represent the dedrifted mappings from each individual to the population-average space. The dedrifted transformations were subsequently used to resample the T1w, T2w, and diffusion tensor images using the appropriate scalar and tensor resampling tools with spline interpolation. The resulting individual images were then averaged to generate the updated population template.

#### Template Generation

Here, we first created an initial fast iterative template using a 240-participant subset of the previously released diffusion data. 5 iterations were run and there was very little difference between iterations 4 and 5 (Figure 2).

**Figure 2:**
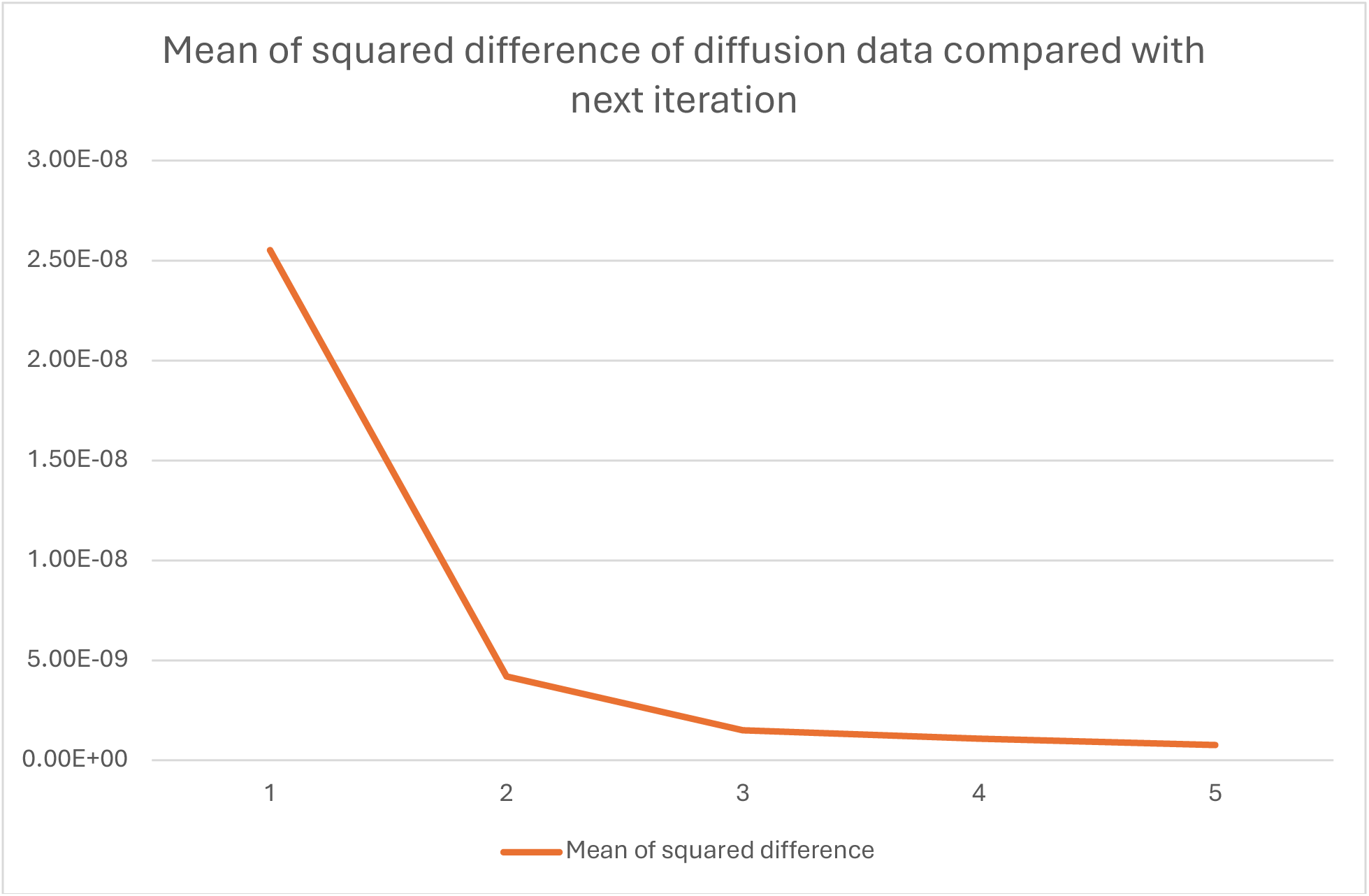
MSE of diffusion data compared with next iteration. The mean squared difference of the tensor template compared with the next iteration was plotted. Virtually no improvement occurs after iteration five, thus indicating convergence of the data.

At this point, we had the reprocessed diffusion data available to us that was described above and we ran two additional iterations with the full dataset of 957 individuals. The mean squared difference between the first and second iterations’ tensor template was 4.059e-10, which was lower than the difference between iterations 5 and 6 of the initial template above, indicating that we had good convergence for the final template.

#### Final Alignment step

All participants from the HCP-YA dataset with sufficient data (n=957) were aligned to the template using identical preprocessing procedures and registration parameters. Corresponding cortical surfaces were generated using the methods from the HCP Pipelines. The resulting subject-level volumetric and surface data were averaged to generate the new population average.

### Inter-individual White matter Fiber Correspondence

To obtain corresponding fibers across individuals for subsequent fiber-preserving tractography matrix averaging, White matter HIerarchical Modelling (WHIM) was used to establish a common set of fiber orientations across participants (Rafipoor et al., 2025). Five iterations of WHIM were performed. At each iteration, subject-level WHIM fitting was performed using the nonlinear transformation obtained from MMORF to sample the diffusion MRI data from native space for each voxel in MMORF group space, with fiber orientations reoriented using the complete Jacobian matrix of the warp and renormalized to unit length (Rafipoor et al., 2025). A common group mask was used for all subjects, generated by averaging the individual T1w brain masks after transformation to MMORF space. The resulting average mask was thresholded at 0.004 (approximately 4/957) to suppress residual mask-related artifacts and subsequently binarized.

For the first iteration, three fibers, corresponding to the number of fibers obtained from bedpostx, were fit to the reprocessed diffusion data using the default WHIM priors. For each subsequent iteration, the WHIM group solution from the preceding iteration was used as the prior for subject-level fitting, thereby progressively refining the correspondence of fiber orientations across individuals. Following subject-level fitting, a group WHIM solution was generated from the current set of individual results using the same group mask. This subject–group fitting procedure was repeated for a total of five iterations, the default set by WHIM.

The group WHIM solution from the fifth iteration, was used as the final template for downstream visualization and correspondence. Following template construction, each individual was fit to the final WHIM template using the subject-level WHIM procedure. These subject-specific fits provided the corresponding individual voxel-wise fiber orientations (fixels) used for subsequent tractography matrix construction.

Fiber orientations were reconstructed within the whole-brain trajectory space at 1.25-mm diffusion resolution using the three posterior fiber populations estimated by WHIM to be viewable by workbench.

### Tractography

**Figure 3:**
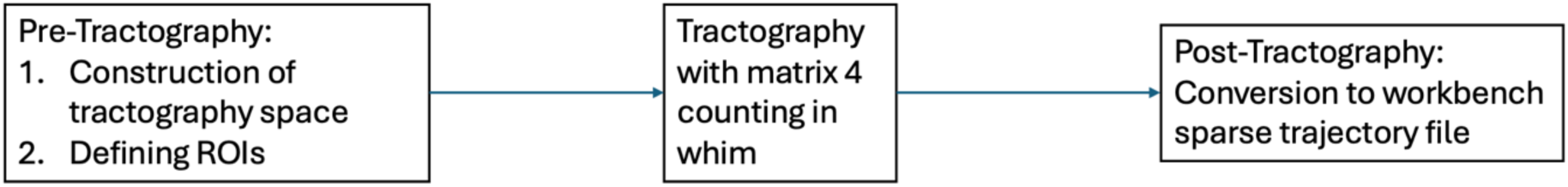
Workflow for tractography. The workflow for generating tractography has three parts. Pre-Tractography needs to be run to construct the tractography space in <u>MM-HCP</u> space and the ROIs used in tractography need to be converted into a format compatible with FSL’s tractography software. Tractography is done using individual white matter voxels as bidirectional seeds. Finally, results of tractography are converted to workbench sparse trajectory files to conserve storage and allow for viewability.

#### Pre-Tractography

##### Construction of the Trajectory Space

A subject-specific trajectory space was generated at the diffusion imaging resolution of 1.25 mm. Anatomical segmentation and cortical surface information were used to construct a volumetric representation of the cortical ribbon and white matter using the native surface in MMORF space. For each hemisphere, signed distance volumes were generated from the native white and pial cortical surfaces. These distance maps were thresholded to obtain binary representations of the regions bounded by the white and pial surfaces. The resulting cortical ribbon was assigned the FreeSurfer cortical labels (3 for the left cortex and 42 for the right cortex), while the white matter regions were assigned the corresponding cerebral white matter labels (2 for the left hemisphere and 41 for the right hemisphere). The hemispheric ribbon volumes were subsequently combined to form a whole-brain ribbon volume at 1.25 mm resolution.

The FreeSurfer white matter parcellation (wmparc) was then used to define the non-cortical components of the trajectory space. Ventricular, cerebrospinal fluid, and choroid plexus structures were explicitly excluded from the valid tracking space. The lateral ventricles, inferior lateral ventricles, third and fourth ventricles, cerebrospinal fluid, and bilateral choroid plexus were excluded. Subcortical gray-matter structures, including the cerebellum, thalamus, basal ganglia, hippocampus, amygdala, and accumbens, were incorporated into the trajectory space using their FreeSurfer labels. Subcortical white matter structures including the ventral diencephalon, the brain stem, and the cerebellar white matter were also incorporated into the trajectory space using their FreeSurfer labels.

The cortical ribbon was subsequently incorporated into the trajectory space using the cortical and cerebral white-matter labels from the ribbon segmentation. Fornix and corpus callosum structures including the posterior, mid-posterior, central, mid-anterior, and anterior corpus callosum subdivisions were additionally incorporated into the trajectory space. These regions were added only where they did not overlap with voxels already assigned to the trajectory space.

The resulting trajectory-space volume was binarized to produce a whole-brain trajectory-space mask. The trajectory-space segmentation was then converted into anatomically defined label volumes using predefined label lookup tables. Separate volumes were generated for whole-brain trajectory space and whole-brain white matter (Figure 4 B). These volumes constituted the subject-specific trajectory-space representation used for subsequent tractography.

**Figure 4:**
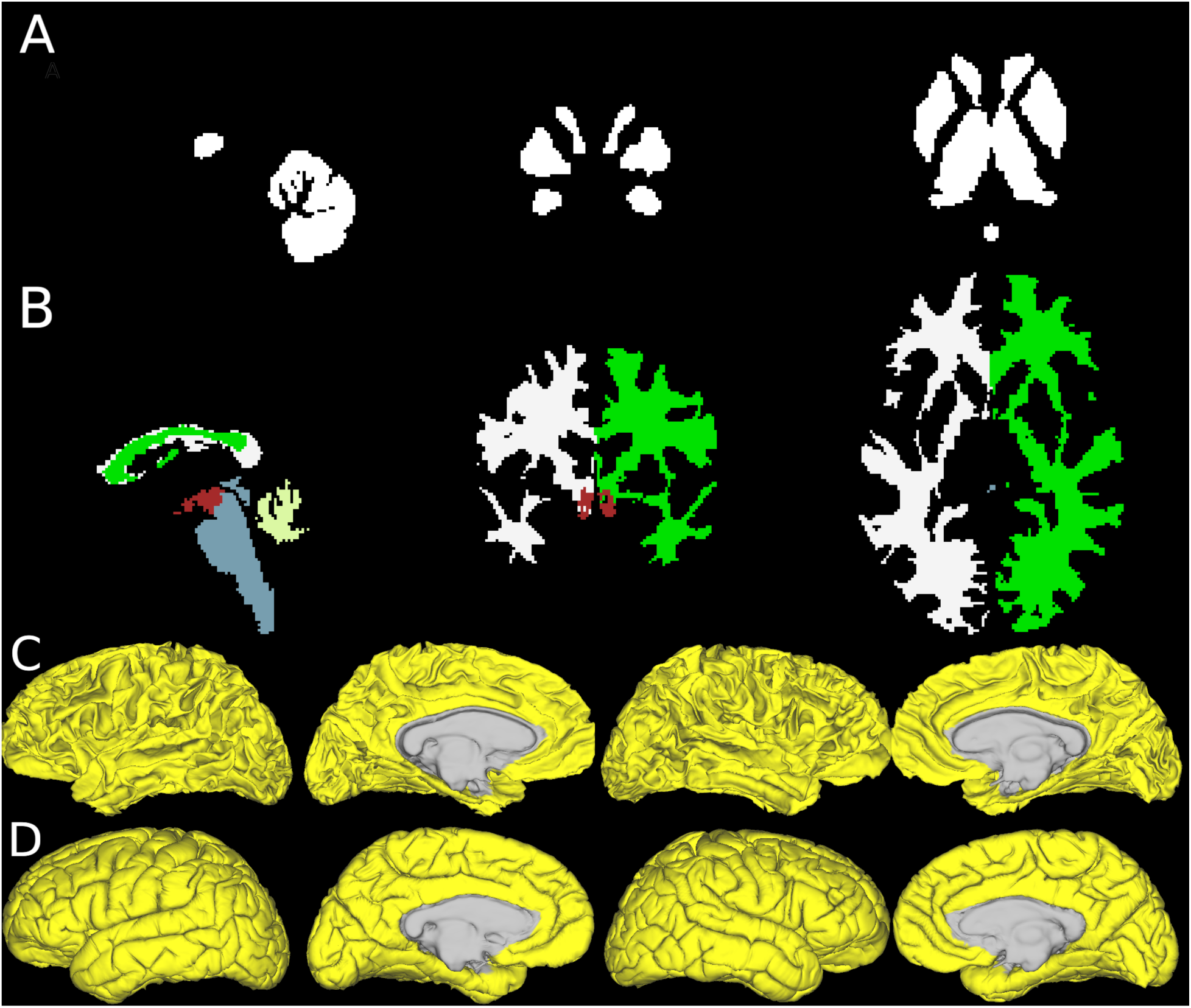
Results of Pre-Tractography. Example result files from Pre-Tractography. The ROI portrayed in A and C contains binarized group average gray matter volumes and MSMAll white surfaces with ROI highlighted in yellow. These will serve as the ROIs for tractography. B shows the sample individual’s whole brain white matter which serves as the seed for tractography. D shows the pial surfaces of the sample individual with yellow being regions where tractography will stop.

#### Definition of Standardized Gray matter and White matter Coordinates

We chose a standard mesh surface resolution of 79212 (“79k”), which has, on average, 1.25mm spacing between its vertices on the midthickness surface of individuals. White and pial surfaces resampled to standard mesh with MSMAll registration were produced in MMORF aligned volume space. To define common regions of interest (ROIs) in each hemisphere across individuals, we used the same atlas ROIs that excluded the medial wall across all individuals (Figure 4 C-D). Thus, each individual had a standard set of cerebral cortical surface vertices for tractography.

Subcortical gray matter was defined using standard volumetric coordinates using the group-level whole-brain subcortical gray-matter atlas at 1.25-mm resolution generated from the mode of individual wmparcs selecting for subcortical gray-matter structures, including the cerebellum, thalamus, basal ganglia, hippocampus, amygdala, and accumbens. This provides a consistent volumetric representation of subcortical gray matter across individuals (Figure 4 A). The cortical and subcortical components were then combined into a CIFTI dense scalar representation, with the cortical components represented on the left and right 79k MSMAll-aligned surfaces and the subcortical component represented volumetrically. This CIFTI file had a total of 231868 grayordinates at 1.25mm average spacing and with each row representing a single grayordinate of the gray matter x white matter x fibers matrix.

The white matter columns were standardized by using the mask used in WHIM analysis and the number of fibers corresponds to the number of fibers obtained from bedpostx.

For tractography, the seed ROI was the individual-specific white matter mask. The individual pial surfaces of both cerebral hemispheres were specified as termination (stop) ROIs, and tracts that exited the trajectory space were also stopped. Streamlines were counted on the white matter surfaces of the cerebral hemispheres and the subcortical gray matter ROIs and streamlines were also not allowed to exit gray matter regions.

#### Probabilistic Tractography

Whole-brain probabilistic tractography was performed for each subject using a modified version of FSL’s probtrackx2_gpu with GPU acceleration. The GPU tractography implementation was extended to support “matrix4” outputs, a matrix in which each row represents a single gray matter location and each column represents the three fibers in a single white matter voxel, with entries containing the count for each gray-matter ID and fiber pair, along with per-voxel distance information, stored in an oxford specific sparse format, and to incorporate fiber-orientation information from the WHIM diffusion model for fiber counting purposes. Because whole-brain tractography produces large connectivity matrices, the previously implemented gray matter to gray matter connectivity matrix (depending on whether seeding began at the gray matter, “matrix1”, or bidirectionally within the white matter, “matrix3”) and “matrix2” gray matter to white matter matrix (without distinctions between multiple fibers in a white matter voxel), representations were converted to sparse matrices using a custom lightweight sparse-matrix class, substantially reducing memory requirements during GPU-based computation.

Tractography was performed using each individual’s BEDPOSTX fiber-orientation distributions in their physical space masked by their whole brain trajectory space. To ensure correspondence between subjects, transformations of seed and streamline coordinates between physical and counting spaces are handled using the forward and inverse MMORF warps.

Individual WHIM fiber orientations in MMORF space were used in the matrix 4 tallying. For each propagated streamline segment, the current fiber direction transformed into counting space was compared with the fiber orientations estimated by WHIM. The WHIM fiber orientation exhibiting the largest absolute dot product with the current streamline direction was identified, thereby assigning each propagated segment to the most directionally corresponding WHIM fiber population.

The whole-brain white-matter volume in MMORF space was used as the seed and waypoint mask, constraining streamline propagation to white-matter tissue. Tractography was seeded throughout the white matter using the matrix 3 seeding strategy to increase sampling of white-matter pathways.

For each white-matter seed voxel, 500 probabilistic samples were generated. Streamline propagation used a step length of 0.3125 mm, corresponding to one quarter of the 1.25-mm voxel resolution, with a maximum of 2,000 steps per streamline. Fiber orientations with a posterior probability below 0.05 were excluded, and loop checking was enabled to prevent anatomically implausible recurrent trajectories. The tractography procedure used forced initial propagation to ensure that streamlines originated from the specified seed locations. Streamlines were terminated upon reaching the gray matter counting regions (cerebral cortical white matter surfaces and subcortical gray matter ROIs), while gray-matter target masks were used to assign reconstructed trajectories to cortical locations. Dense connectivity matrices were generated including gray matter x gray matter counts and distances (matrix 3) and gray matter x white matter x fibers + distance (matrix4).

#### Post processing

For each subject, the individual Matrix 4 outputs were subsequently converted into Connectome Workbench sparse matrix format (fdt_matrix4.trajTEMP.wbsparse). Specifically, the matrix4 output was provided together with the subject-specific tract-space coordinates and the individual’s Diffusion.Whim fiber orientation file. Cortical seed locations were defined using the subject’s 79k_fs_LR Gray.dscalar.nii file, with the COLUMN dimension specifying the seed dimension. This conversion provided a common representation of the individual tractography results in the group-level fiber space, allowing the subject-specific connectivity information to be subsequently combined or compared across individuals.

### Non-negative Matrix Factorization

We averaged across individuals to assess the typical pattern of structural connectivity in the young adult brain. We used a probabilistic streamline threshold of 3, determined by the elbow of fiber counts, to minimize the RAM requirements for NMF (as most non-zero entries in the sparse matrix had very few streamlines on average) (Figure 5). Here, we adapted a very fast sparse NMF algorithm based on RcppML (DeBruine et al., 2024) in R. We addressed an int_32 overflow issue with the R based sparse matrix indexes used in the original algorithm by converting the underlying data structure and matrix operations to eigen library based sparse matrix (Guennebaud et al., 2010) in C++. We also halved the RAM requirements by altering the method of predicting **W** based on **H** to no longer store the transpose of the original matrix, without affecting the processing speed by precomputing the **W^T^H** in a parallelized column major favorable manner.

**Figure 5:**
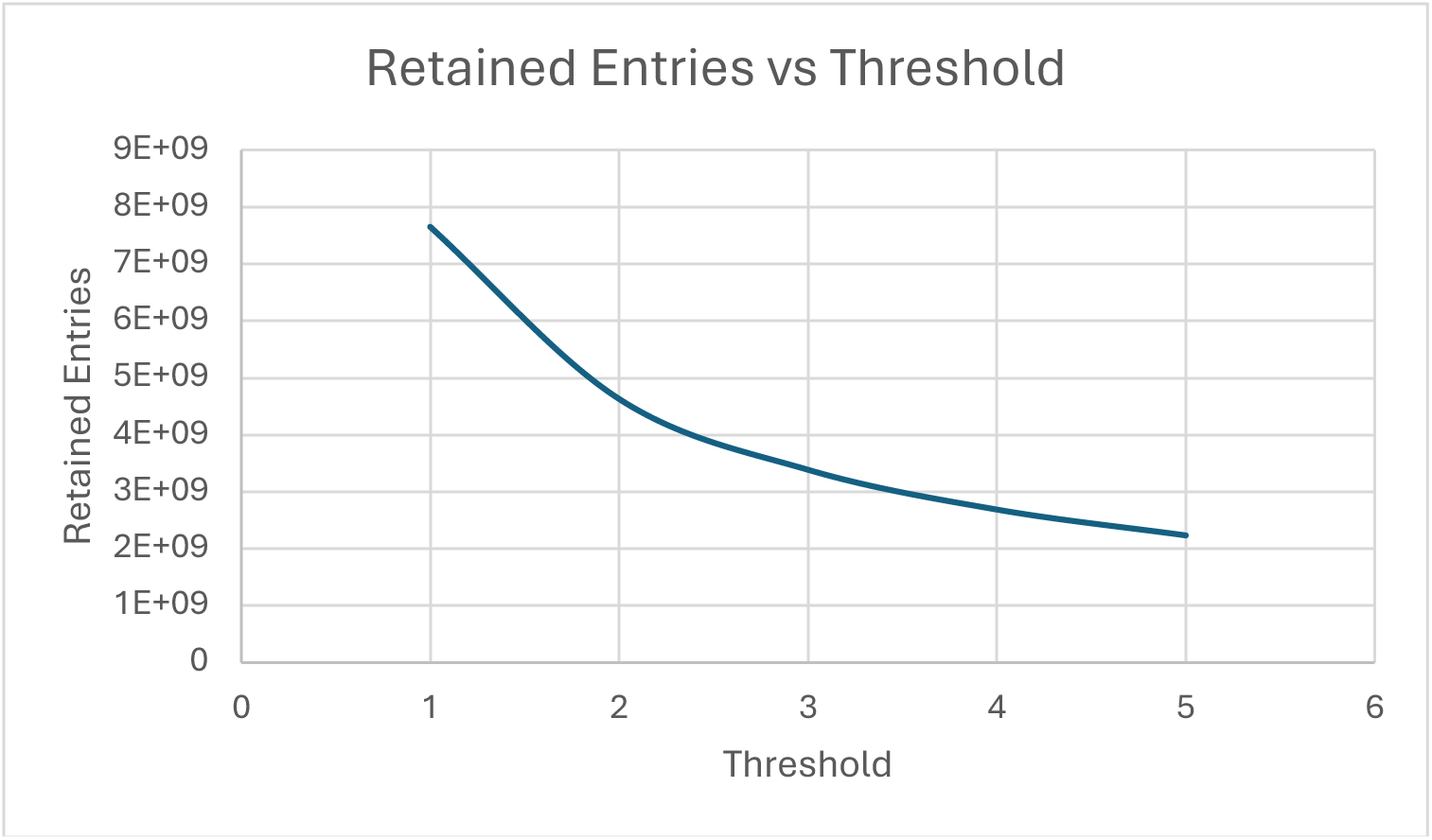
Retained entry against threshold. The number of entries retained is plotted against the threshold that is used. The threshold is applied to per fiber counts greater than or equal to the threshold. The elbow point is threshold=3.

#### NMF Stability and Consensus Analysis

To assess the stability and reproducibility of the non-negative matrix factorization (NMF) solutions, an NMF(ICA)SSO (Himberg & Hyvarinen, 2003)-based stability analysis was performed across repeated NMF runs. Each NMF repetition produced a factorization consisting of a basis matrix (W), coefficient matrix (H), and component-specific strength parameter (d). The resulting components from all valid NMF repetitions were pooled and treated as individual estimates for subsequent clustering.

For each pair of NMF component estimates, similarity was quantified using the Pearson correlation between their corresponding (W) vectors. Negative component correlations were set to zero. The resulting component-by-component similarity matrix was converted to a dissimilarity matrix using the square-root similarity-to-dissimilarity transformation implemented in ICASSO. The individual component estimates were then hierarchically clustered using agglomerative clustering with average linkage.

The number of clusters was fixed to the number of NMF components in the original factorization. Cluster stability was evaluated using the ICASSO stability index (I_q), which quantifies the reproducibility of components across the independent NMF repetitions. The ICASSO projection procedure was additionally applied using canonical correlation analysis to obtain a low-dimensional representation of the component estimates and facilitate identification of representative components.

For each stability cluster, a centrotype was selected as the representative NMF component. The (W), (H), and (d) parameters associated with these centrotypes were extracted to form a consensus NMF solution. The resulting components were subsequently ordered according to descending component strength (d). Thus, the consensus (W) and (H) matrices represent the most representative and reproducible components across the repeated NMF solutions rather than the result of a single NMF initialization.

For each decomposition rank (K = 25, 50, and 75), 100 random NMF initializations were performed on the group-average connectome of 100 individuals using the NMFSSO approach. The resulting solutions were used to derive the final initializing W for a subsequent final NMF run, which is more reproduceable than any single random NMF run.

## Results

### Group Average Template and novel MM-HCP standard space

To obtain volumetric alignment between individuals, the new population-specific template was generated using the MMORF registration framework to represent the young adult population while maintaining the ACPC orientation convention as described above. The final template and standard space was generated from 957 individuals based on T1w, T2w, and diffusion tensor images. Group averages of the T1w and T2w images exhibited sharp subcortical gray and white matter anatomical boundaries (Figure 6 A, B). The group average diffusion tensor shows sharp white matter tracts colored according to the principle eigen vector with the color intensity determined by the fractional anisotropy (Figure 6 C). Cortical surfaces show the consistent cortical folds across individuals (Figure 6 D). These data illustrate excellent inter-individual correspondence.

**Figure 6:**
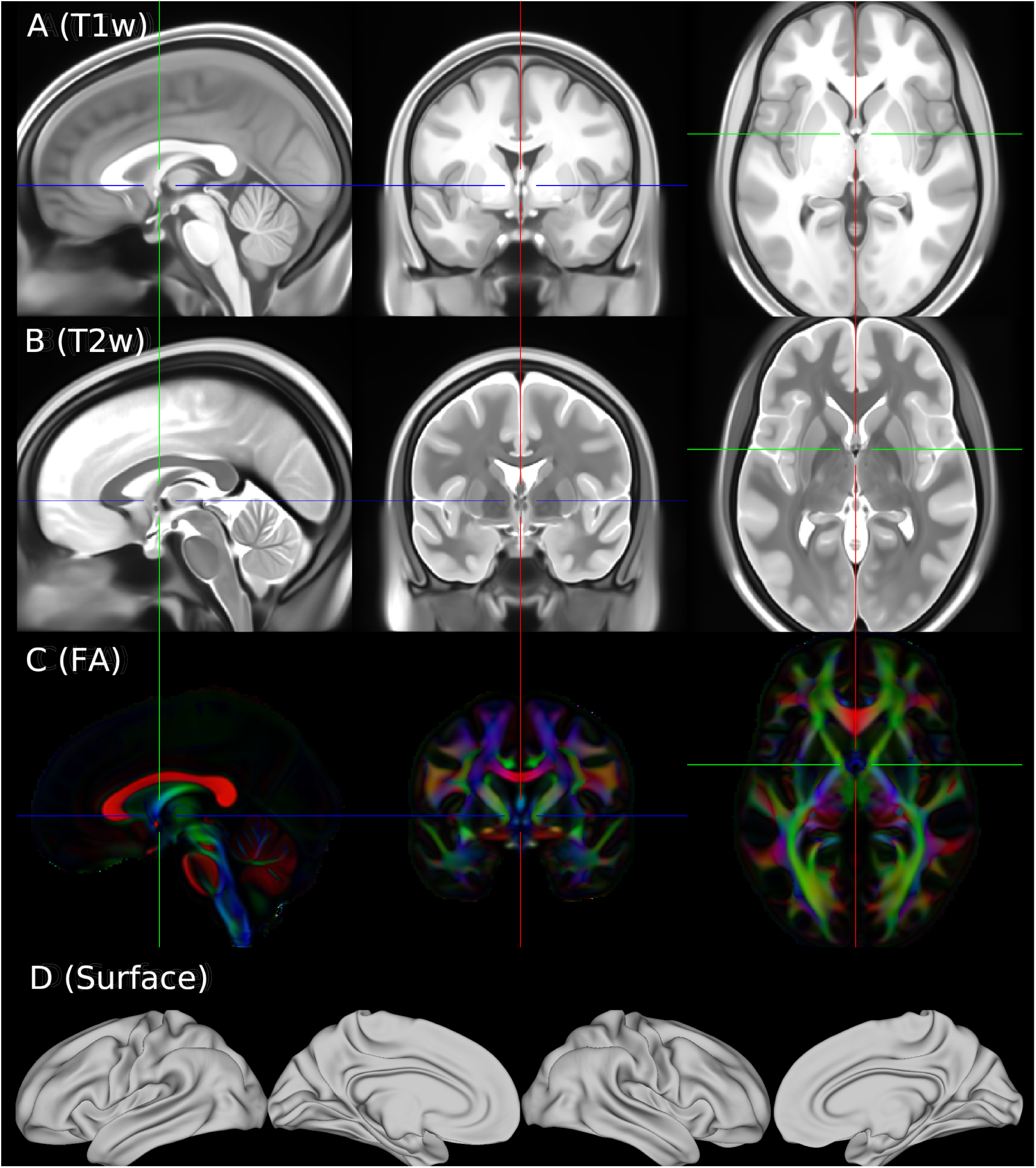
Group average of 957 HCP-YA individuals in MM-HCP space. The group average aligned with the template in MM-HCP space is shown with first row T1w, second row T2w, third row colored FA map, fourth row average white matter surface. The colored FA map is masked using the mask generated from WHIM for better visualization.

Furthermore, to obtain fiber correspondence, a new WHIM template was created in MM-HCP space to align fibers across individuals. Figure 7 shows spatially organized fiber orientations across sagittal and coronal views, demonstrated by the centrum semi-ovale with fiber 1, 2, 3 encoded as red, green, and blue respectively (Figure 7 A). When compared with an exemplar individual with the same color encoding (Figure 7 B), the first, second, and third fibers are directionally consistent. The resulting template provides a group level representation of fiber orientations, allowing cross-individual fiber comparisons.

**Figure 7.**
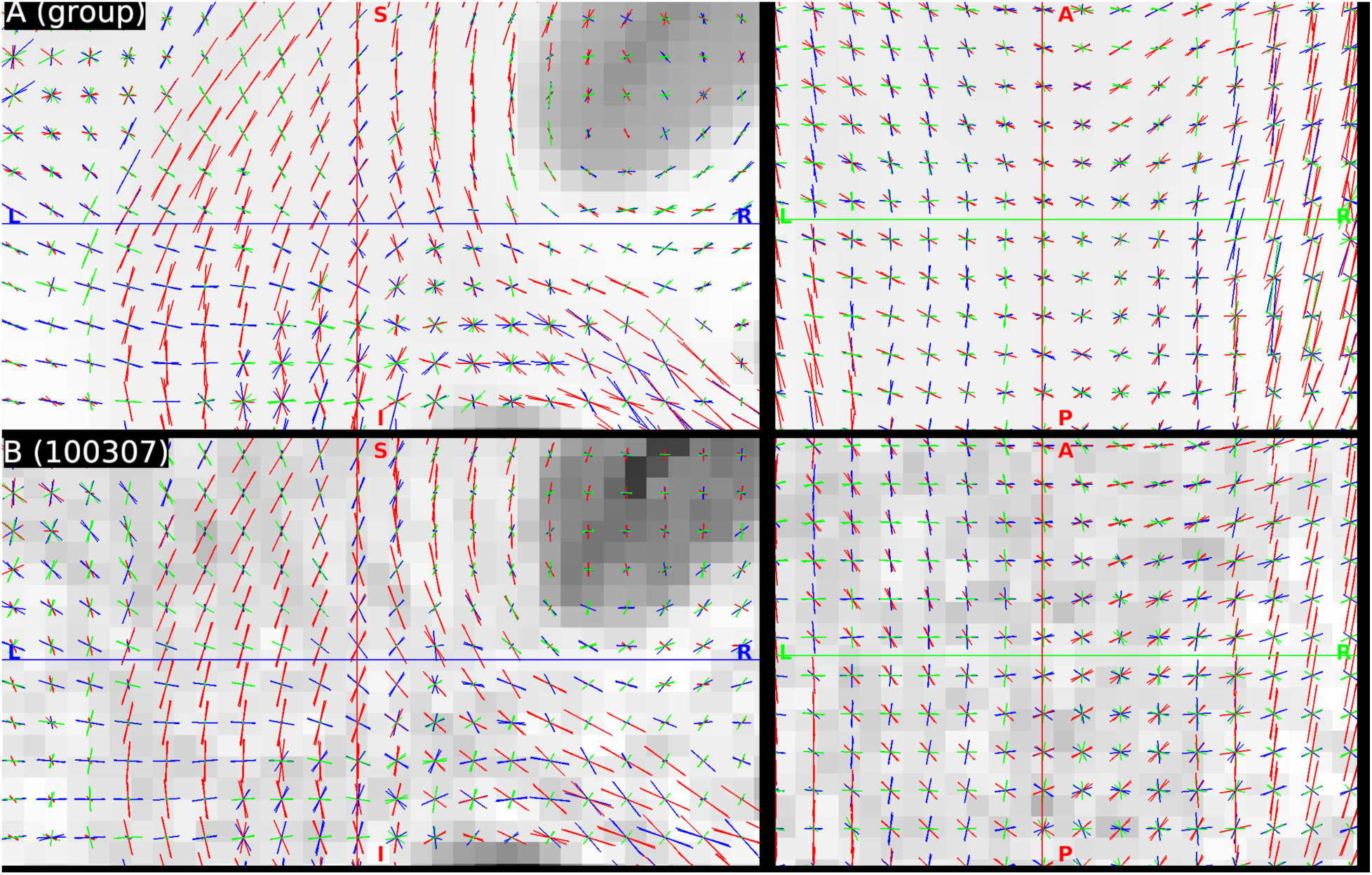
Whim template and individual-subject fiber orientations in MM-HCP space. The WHIM template was created in MM-HCP space with the average T1w brain mask thresholded to 0.004 and after 5 iterations of template generation. The individual map is generated from an extra iteration on an individual (HCP subject 100307). The template fibers and individual fibers are overlaid on top of the group average brain extracted T1w image (A) and individual T1w image (B) with fiber 1,2,3 as RGB respectively.

### NMF of Group Average Tractography

To assess the reproducibility of tractography derived NMF components, individual tractography results were averaged within each of two groups that are subsets of 475P and 475V (Yang et al., 2025) with each group containing 100 individuals. The NMFSSO approach was applied separately to each group’s group-averaged trajectory matrix, using 25, 50, 75 components respectively, with reproducibility plot shown in in Figure 8.

**Figure 8:**
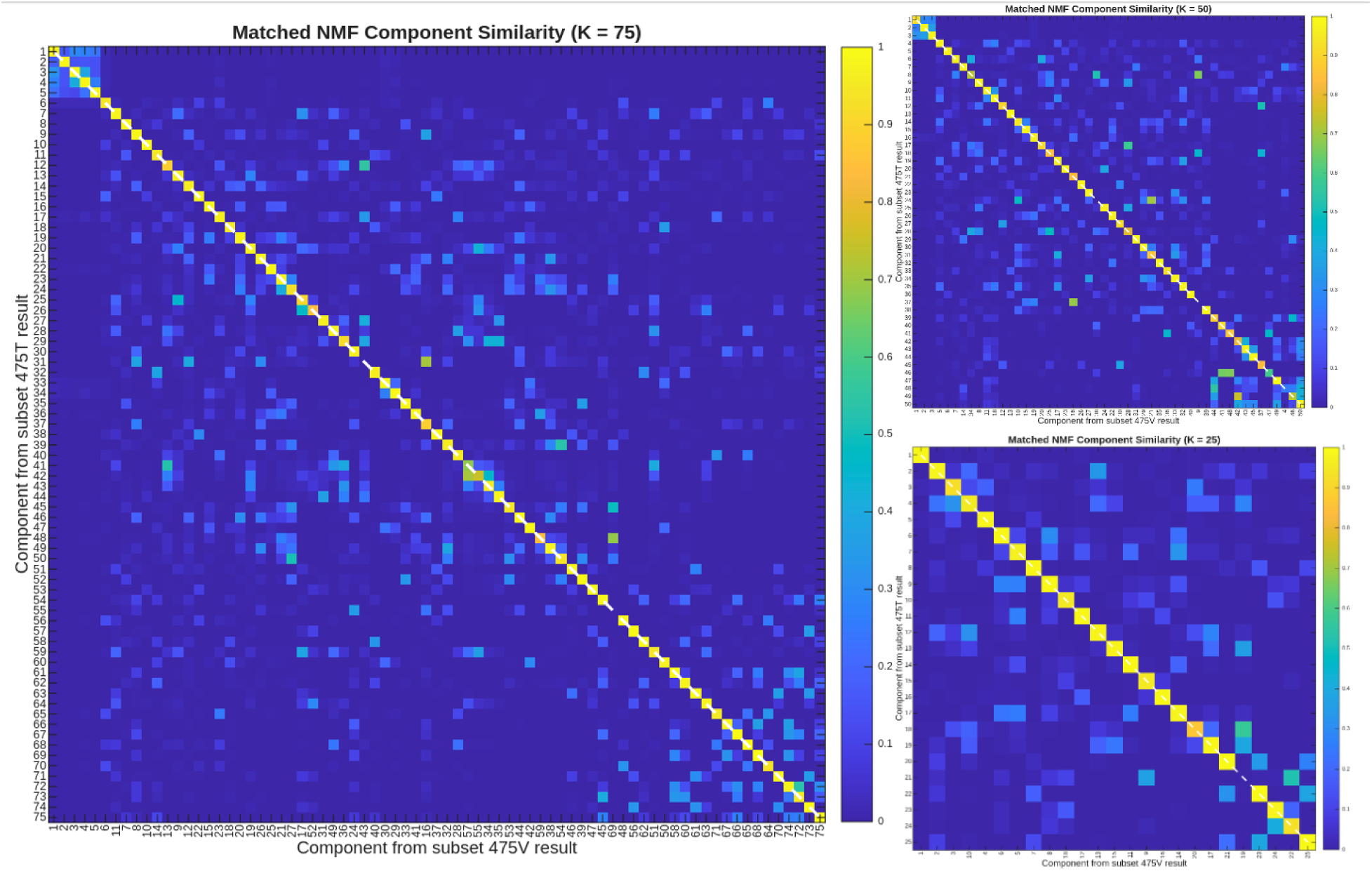
Reproducibility plot of 25, 50, and 75 NMF component decompositions. These were performed on group average tractography matrices from two independent subsets of 100 individuals from the 475P and 475V groups using the NMFSSO method described above. Components are matched based on cosine similarity. The mean matched cosine similarities were 0.93, 0.88, and 0.94 for K = 25, 50, and 75, respectively.

Figure 8 shows matched exemplar components (left and right hemisphere portions of the Inferior Longitudinal Fasciculus; ILF) from each group from the d=75 component decomposition, paired by their best match across groups using cosine similarity. White matter trajectories are shown using the chosen (of the available 3) WHIM fiber orientations (green representing anterior to posterior fibers, red representing medial to lateral fibers, and blue representing superior to inferior fibers) displayed over group average cortical surfaces, with the fiber trajectories displayed in front of the surface even though they are actually interiorly located. The opacity of each fiber is determined by the number of streamlines that chose that fiber. Cortical surface maps of grey matter projection territories are the proportion of the total decomposed streamlines at each vertex location (0-100%). This visualization approach reduces the effect of the well-known gyral bias phenomenon (Van Essen et al., 2014). Matched (across the independent decompositions) NMF components are shown for superior and inferior ILF subdivisions in the left and right hemispheres. Cosine similarity values were 0.995 and 0.996 respectively for the superior and inferior ILF subdivisions in the left hemisphere and 0.994 and 0.996 respectively for these subdivisions in the right hemisphere.

**Figure 9:**
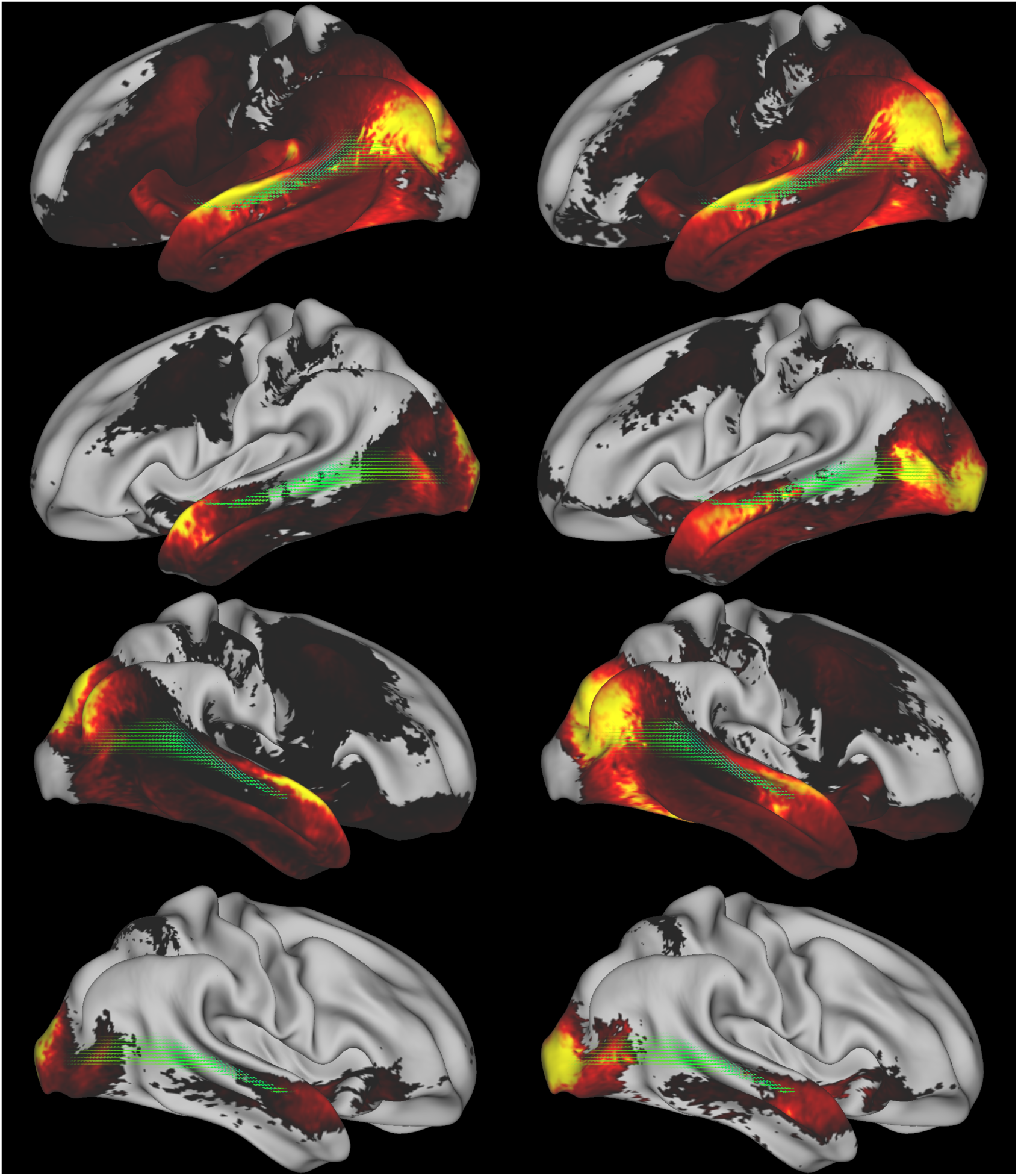
Results of NMF of the group average. Sample components shown here are: left (100 individual subset from 475V) component 61, 69, 63, and 74, right (100 individual subset from 475P) component 58, 68, 61, 73. Fibers are colored green (anterior to posterior), red (medial to lateral), blue (superior to inferior), and their opacity is determined by the number of streamlines choosing them. Grey matter projection territories are colored by the proportion of the decomposed streamlines at each vertex represented by each component.

## Discussion

In this study, we implemented methods for data-driven decomposition of precisely aligned, fiber-orientation-preserving probabilistic tractography connectomes using non-negative matrix factorization (NMF). Several prior methodological advances enabled this approach, including the development of the MMORF algorithm for multi-modal nonlinear volumetric alignment (Lange et al., 2024), the WHIM algorithm for aligning fiber directions across individuals (Rafipoor et al., 2025), sparse NMF algorithms capable of handling extremely large sparse datasets (DeBruine et al., 2024), and better computer hardware technologies including increases in RAM capacity, storage capacity, and increases in CPU and GPU performance. Methodological contributions of this study include an HCP Pipeline for MMORF, improvements to FSL’s probtrackx, including implementation of matrix4 tractography on the GPU, WHIM-based matrix4 counting, and improving memory efficiency to support high-resolution whole-brain analyses, an HCP tractography pipeline, improving the RAM efficiency of a sparse NMF algorithm to accommodate these extremely large datasets, and adaption of ICASSO to NMFSSO. Additionally, we implemented multiple improvements to Connectome Workbench and its command line utilities to support this work. Together, with these algorithmic and hardware improvements, we demonstrated that it is possible to conduct whole brain fiber-preserving probabilistic tracking at high resolution and that NMF can produce reproducible high-resolution group average decompositions of the entire adult human brain. We also produced a new unbiased MM-HCP standard volume space from HCP young adults that adheres to ACPC orientation conventions.

The tools developed by this study enable data-driven decomposition of probabilistic dense connectomes while preserving complex fiber architectures into white matter tracts and gray matter projection territories. This approach represents an alternative to ROI-based tract definitions (akin to the model-free decompositions of fMRI data vs seed-based connectivity analyses). Here, we showed that such decompositions are reproducible and reflect known white matter tracts and they could be projected back to individuals using non-negative dual regression approaches (Filippini et al., 2009; Thompson et al., 2020). Such methods would enable fully data-driven imaging-derived phenotypes (IDPs) based on white matter tracts, without requiring ROI-based definitions of such tracts and also avoiding the limitations of existing model-free voxelwise methods like TBSS (Smith et al., 2006).

Several directions remain to be explored. First, the optimal number of components should be determined using the full cohort of 957 individuals. Potential approaches include identifying a knee point in the relationship between model rank and reconstruction error or identifying local minima in reconstruction error. Second, once we determine the optimal number of components, we need to identify the tracts neuroanatomically (including relation to ROI-based atlases like XTRACT (Warrington et al., 2020)) and relate them to atlases of brain areas and functional networks that are under development.

There are several limitations to this approach. First, despite the many methodological advancements enabling feasibility, this study has still been a computationally expensive effort, requiring access to a center for high-performance computation (CHPC) at our institution. Computational requirements would be reduced for more typical diffusion resolutions such as 1.5mm or 2mm and advances in computer hardware are likely to continue for some time (though advances in high-field acquisition technologies could bring such analyses at state of the art acquisition resolutions out of reach again due to further gains in spatial resolution (Feinberg et al., 2023)). That said, diffusion tractography is a tremendously resolution starved problem, with several hundred thousand axons passing through a square mm cross-section of white matter (Aboitiz et al., 1992). Second, these approaches still cope with inherent limitations of tractography, such as gyral (or anti-sulcal) biases, biases towards the geometrically easiest paths, the inability to distinguish crossing fibers from axonal bifurcations and potential for scrambling tracts within deep fascicles. Importantly, though, these issues are most important for area-to-area connectomes, whereas here our goal was to reconstruct fiber fascicles and their entire gray matter projection territories.

Overall, this study represents a concerted effort to extract more value from the HCP’s diffusion tractography structural connectivity data to enable greater parity between it and the thus-far more successful functional connectivity data. We believe a data driven atlas of white matter tracts and their gray matter projection territories will enable useful data-derived tract-based imaging-derived phenotype (IDP) analyses. Further, many studies may benefit from the unbiased, better aligned, young-adult-based HCP-derived standard space, including those interested in the globular gray matter subcortical nuclei.

## Acknowledgements

The authors wish to acknowledge funding from NIH NIA R01AG092088 and Matthew Webster and Jesper Andersson for helpful discussions.

